# Host interactions of novel *Crassvirales* species belonging to multiple families infecting bacterial host, *Bacteroides cellulosilyticus* WH2

**DOI:** 10.1101/2023.03.05.531146

**Authors:** Bhavya Papudeshi, Alejandro A. Vega, Cole Souza, Sarah K. Giles, Vijini Mallawaarachchi, Michael J. Roach, Michelle An, Nicole Jacobson, Katelyn McNair, Maria Fernanda Mora, Karina Pastrana, Lance Boling, Christopher Leigh, Clarice Harker, Will S. Plewa, Susanna R. Grigson, George Bouras, Przemysław Decewicz, Antoni Luque, Lindsay Droit, Scott A. Handley, David Wang, Anca M. Segall, Elizabeth A. Dinsdale, Robert A. Edwards

## Abstract

Bacteroides, the prominent bacteria in the human gut, play a crucial role in degrading complex polysaccharides. Their abundance is influenced by phages belonging to the *Crassvirales* order. Despite identifying over 600 *Crassvirales* genomes computationally, only few have been successfully isolated. Continued efforts in isolation of more *Crassvirales* genomes can provide insights into phage-host-evolution and infection mechanisms. We focused on wastewater samples, as potential sources of phages infecting various *Bacteroides* hosts. Sequencing, assembly, and characterization of isolated phages revealed 14 complete genomes belonging to three novel *Crassvirales* species infecting *Bacteroides cellulosilyticus* WH2. These species, *Kehishuvirus* sp. ‘tikkala’ strain Bc01, *Kolpuevirus* sp. ‘frurule’ strain Bc03, and ‘Rudgehvirus jaberico’ strain Bc11, spanned two families, and three genera, displaying a broad range of virion productions. Upon testing all successfully cultured *Crassvirales* species and their respective bacterial hosts, we discovered that they do not exhibit co-evolutionary patterns with their bacterial hosts. Furthermore, we observed variations in gene similarity, with greater shared similarity observed within genera. However, despite belonging to different genera, the three novel species shared a unique structural gene that encodes the tail spike protein. When investigating the relationship between this gene and host interaction, we discovered evidence of purifying selection, indicating its functional importance. Moreover, our analysis demonstrated that this tail spike protein binds to the TonB-dependent receptors present on the bacterial host surface. Combining these observations, our findings provide insights into phage-host interactions and present three *Crassvirales* species as an ideal system for controlled infectivity experiments on one of the most dominant members of the human enteric virome.

**Impact statement:** Bacteriophages play a crucial role in shaping microbial communities within the human gut. Among the most dominant bacteriophages in the human gut microbiome are *Crassvirales* phages, which infect Bacteroides. Despite being widely distributed, only a few *Crassvirales* genomes have been isolated, leading to a limited understanding of their biology, ecology, and evolution. This study isolated and characterized three novel *Crassvirales* genomes belonging to two different families, and three genera, but infecting one bacterial host, *Bacteroides cellulosilyticus* WH2. Notably, the observation confirmed the phages are not co-evolving with their bacterial hosts, rather have a shared ability to exploit similar features in their bacterial host. Additionally, the identification of a critical viral protein undergoing purifying selection and interacting with the bacterial receptors opens doors to targeted therapies against bacterial infections. Given Bacteroides role in polysaccharide degradation in the human gut, our findings advance our understanding of the phage-host interactions and could have important implications for the development of phage-based therapies. These discoveries may hold implications for improving gut health and metabolism to support overall well-being.

**Data summary:** The genomes used in this research are available on Sequence Read Archive (SRA) within the project, PRJNA737576. *Bacteroides cellulosilyticus* WH2, *Kehishuvirus* sp. ‘tikkala’ strain Bc01, *Kolpuevirus sp. ‘*frurule’ strain Bc03, and ‘Rudgehvirus jaberico’ strain Bc11 are all available on GenBank with accessions NZ_CP072251.1 (*B. cellulosilyticus* WH2), QQ198717 (Bc01), QQ198718 (Bc03), and QQ198719 (Bc11), and we are working on making the strains available through ATCC. The 3D protein structures for the three *Crassvirales* genomes are available to download at doi.org/10.25451/flinders.21946034.

## Introduction

The intricate relationship between gut microbiomes and human health is characterized by the diverse microbial communities that help with digestion, regulate the immune system, and alter brain function (1–3). Metagenomics, a culture-independent technique is used to capture microbial diversity in a sample (4,5), has transformed our understanding of bacteria and the corresponding bacteriophages in the environment (6–9). These metagenomic datasets have revealed a correlation between bacterial and bacteriophage populations, which suggests bacteriophages play a role in modulating bacterial populations (10,11). In particular, the human gut microbiome exhibits varying bacterial densities, including a high abundance of Bacteroidota (formerly Bacteroidetes) (12–14), which are shaped by the phages belonging to the *Crassvirales* order. These dsDNA bacteriophages have a podovirus-like morphology, genomes ranging between 100 and 200 kb, and conserved gene order (15–17). They are widespread, constituting a stable component of an individual’s microbiome, and do not appear to be associated with human health or disease states (16,18).

The first phage within *Crassvirales* order was computationally discovered by cross-assembly of DNA sequence reads from human gut microbiome samples (19). Since, nearly 600 *Crassvirales* genomes have been identified computationally, leading to the International Committee of Taxonomy of Viruses (ICTV) formally classifying the *Crassvirales* order into four families, ten subfamilies, 42 new genera, and 72 new species (17,20). The classification relied on phylogenetic analysis of conserved structural genes, including the major capsid protein (MCP), terminase large subunit (*terL*), and portal protein (portal). Additionally, the average nucleotide identity (ANI) species cutoff was set to 95 % identity over 85 % genome coverage.

The identification of numerous *Crassvirales* genomes has advanced our understanding of this viral order. Similar to other phages, these genomes contain three discernible regions encoding for 1) structural proteins involved in producing the capsid and tail genes, 2) transcription proteins and 3) replication proteins crucial for successful phage replication in different infection stages (15). Gene homology analysis showed that majority of the genes are highly variable when compared with other genomes from this order. Comparative genomics further displayed unique biological characteristics of *Crassvirales* species (21,22), including switching DNA polymerases, alternative coding strategies (23–26), and the variable intron density across lineages (15,26). Overall, functional annotation of *Crassvirales* genomes remain challenging, with majority of the genes annotated as hypothetical proteins lacking known biological function, with little to no similarity with sequences in reference databases. These challenges can be addressed through experimental approaches that can help elucidate the functions of these uncharacterized genes or proteins.

The first step in experimental approaches is phage isolation, which requires the knowledge of their host species, and the ability to culture them. This has led to only four successful isolates obtained so far, including *Kehishuvirus primarius* (crAss001) infecting *Bacteroides intestinalis* APC919/174 (27), *Wulfhauvirus bangladeshii* DAC15 and DAC17 from wastewater effluent infecting *Bacteroides thetaiotaomicron* VPI-5482 (28),and *Jahgtovirus secundus* (crAss002) infecting *Bacteroides xylanisolvens* APCS1/XY(29). All these isolates exhibited host specialist morphotypes that can be maintained in the continuous host culture, but none possess lysogeny-related genes (27,30). The proposed mechanism of persistence includes where the bacterial host cycles between sensitive and resistant states through altering the genes encoding surface transporters and capsular polysaccharide structures (CPS) on the bacterial surface (30,31). Further to improve the annotations, cryogenic-electron microscopy of *K. primarius* provided functional assignments to the virion proteins, and an insight into the infection mechanism, revealing how the capsid and tail store cargo proteins aid in initial host infection (32). Continued efforts in isolation of more *Crassvirales* genomes can provide insights into phage-host-evolution, comprehensive protein annotation and elucidation of infection mechanisms.

Here we present the successful isolation of 14 *Crassvirales* isolates from wastewater, that include three novel species, belonging to families *Steigviridae* and *Intestiviridae.* Remarkably, all these isolates infect the same host, *Bacteroides cellulosilyticus* WH2. We investigate the genes playing a role in host interaction, providing insights into the evolution of these dominant phages, and how their interactions shape the gut microbiome.

## Results

### Search for *Crassvirales* phages

We obtained a total of 41 phages from wastewater infecting four different *Bacteroides* species, *B. cellulosilyticus* WH2, *B. fragilis* NCTC 9343, *B. stercoris* CC31F, and *B. uniformis* ATCC 8492. The phages were sequenced using Oxford Nanopore or Illumina MiSeq platforms, and the resulting sequences were assembled. We performed BLASTN searches of the assembled phages against the non-redundant (nr/nt) NCBI database for taxonomic assignment. As a result, we identified 14 phages infecting *B. cellulosilyticus* WH2 belong to *Crassvirales* order. Each of these phages were labelled with code ranging from Bc01 to Bc14.

### Isolation and taxonomic classification of *Crassvirales* isolates

*Crassvirales* isolates formed distinct clear circular plaques with a uniform diameter of 1 mm on soft agar overlays. We performed shotgun sequencing on the 14 isolates phages, with Bc01 to Bc03, Bc05 to Bc11 sequenced on Oxford Nanopore platform, and Bc01 to Bc08, Bc12 to Bc14 sequenced on the Illumina platform. The assemblies produced multiple contigs, and our selection criteria to identify complete phage genomes was based on presence of viral genes, highest read coverage and unitigs (high quality contig) size of approximately 100 kb (Table 1). This resulted in complete genomes for each of the 14 phages, that were polished with Illumina reads correcting for substitution, insertion, and deletion errors.

**Table 1:** Taxonomic classification of the 14 *Crassvirales* genomes isolated from wastewater infecting *Bacteroides cellulosilyticus* WH2.

| Phage isolate | Sequencing platform | Genome length | Taxonomy | Biosample ID |
| --- | --- | --- | --- | --- |
| **Bc01** | MinION, MiSeq | 100,722 | *Kehishuvirus* sp. ‘tikkala’ strain Bc01 | SAMN20326212 |
| **Bc02** | MinION, MiSeq | 98,905 | *Kolpuevirus* sp. ‘frurule’ strain Bc02 | SAMN20326213 |
| **Bc03** | MinION, MiSeq | 99,379 | *Kolpuevirus* sp. ‘frurule’ strain Bc03 | SAMN20326214 |
| **Bc04** | MiSeq | 99,033 | *Kolpuevirus* sp. ‘frurule’ strain Bc04 | SAMN29929441 |
| **Bc05** | MinION, MiSeq | 97,832 | *Kolpuevirus* sp. ‘frurule’ strain Bc05 | SAMN20326216 |
| **Bc06** | MinION, MiSeq | 99,845 | *Kolpuevirus* sp. ‘frurule’ strain Bc06 | SAMN20326217 |
| **Bc07** | MinION, MiSeq | 98,518 | *Kolpuevirus* sp. ‘frurule’ strain Bc07 | SAMN20326218 |
| **Bc08** | MinION, MiSeq | 98,067 | *Kolpuevirus* sp. ‘frurule’ strain Bc08 | SAMN20326219 |
| **Bc09** | MinION | 98,788 | *Kolpuevirus* sp. ‘frurule’ strain Bc09 | SAMN20326220 |
| **Bc10** | MinION | 96,329 | *Kolpuevirus* sp. ‘frurule’ strain Bc10 | SAMN20326221 |
| **Bc11** | MinION | 90,458 | ‘Rudgehvirus jaberico’ strain Bc11 | SAMN20326222 |
| **Bc12** | MiSeq | 96,952 | *Kolpuevirus* sp. ‘frurule’ strain Bc12 | SAMN29929442 |
| **Bc13** | MiSeq | 90,716 | ‘Rudgehvirus jaberico’ strain Bc13 | SAMN29929443 |
| **Bc14** | MiSeq | 98,803 | *Kehishuvirus* sp. ‘tikkala’ strain Bc14 | SAMN29929444 |

For taxonomic classification of these isolates, we applied the ICTV report guidelines for defining taxonomy within *Crassvirales* order. Phylogenetic clustering of the conserved portal gene and average nucleotide identity (ANI) species cutoff (95 % identity over 85 % genome coverage) identified three distinct clusters (Figure 1A). We selected the highest confidence genomes: Bc01, Bc03, and Bc11 from each cluster. These three isolates were compared against all known *Crassvirales* genomes through phylogenetic clustering of conserved genes, major capsid protein (MCP), portal, terminase large subunits (*terL*) genes to determine that Bc01 and Bc03 clusters belong to the *Steigviridae* family, and Bc11 to the *Intestiviridae* family (Figure 1B, Figure S1). Confirmation of the genus assignment was obtained through ANI and shared protein information, which identified Bc01 to *Kehishuvirus*, Bc03 to *Kolpuevirus*, and Bc11 to a novel genus group that we propose to name ‘Rudgehvirus*’*.

**Figure 1:**
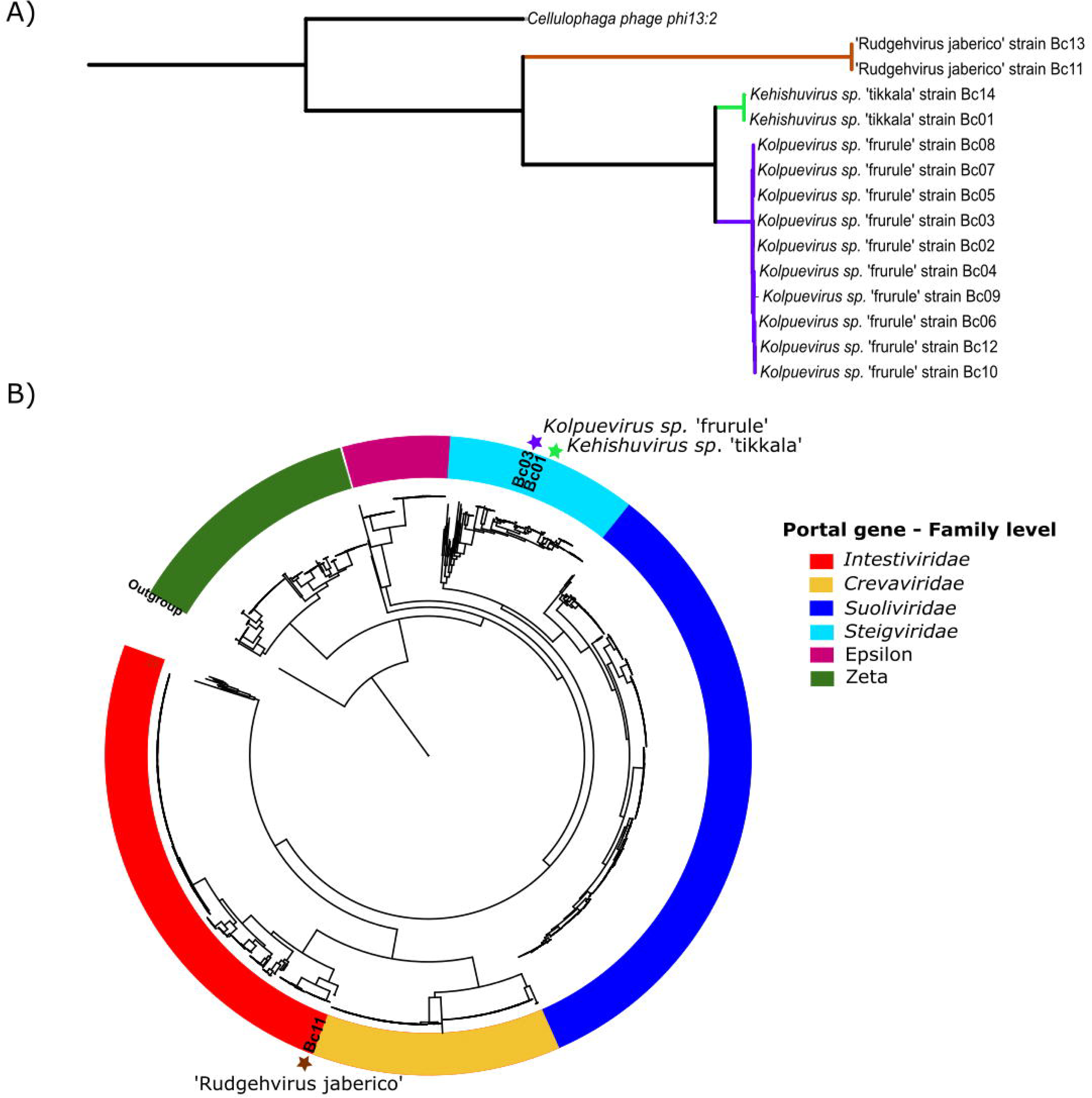
Phylogenetic tree constructed using the portal protein using JTT model, CAT approximation with 20 rate categories and outgroup set to Cellulophaga phage phi13:2 A) Phylogenetic tree of the 14 *Crassvirales* isolates with the branches color-coded to represent the three species, *Kehishuvirus* sp.in light green, *Kolpuevirus* sp. in purple, and ‘Rudgehvirus’ in brown. B) Clustering of all known *Crassvirales* genomes confirming that isolate *Kehishuvirus* sp. ‘tikkala’ strain Bc01 and *Kolpuevirus* sp. ‘frurule’ strain Bc03 belong to the family Steigviridae (cyan), and ‘Rudgehvirus jaberico’ strain Bc11 to Intestiviridae (red).

All three isolates represent novel species exhibited less than 95% identity and 85% coverage to any known *Crassvirales* genomes. Bc01 is most similar to the reference genome *Kehishuvirus primarius* (Genbank ID: MH675552) with 95.5% identity across 79.1 % genome coverage. Bc03 aligns with *Kolpuevirus hominis* (Genbank ID: MT774391) with 82.8 % identified across 53.73 % query coverage. Bc11 aligns with the reference genome *Jahgtovirus intestinalis* (Genbank ID: OGOL01000109) with 74.7 % identity across only 9.9 % query coverage. We proposed names for these novel species as *Kehishuvirus* sp. ‘tikkala’ strain Bc01, *Kolpuevirus* sp ‘frurule’ strain Bc03, and ‘Rudgehvirus jaberico’ strain Bc11.

### Genome characteristics of the novel *Crassvirales* species

*Kehishuvirus* sp. ‘tikkala’ strain Bc01 is 100,841 bp, with 104 proteins, 24 tRNAs and GC content of 35.09 % (Table 2) which is lower than the bacterial host GC content of 42.8 %. These isolates formed clear, uniform circular spot plaques approximately 1 mm in diameter, forming 9.3*10^9^ PFU/mL (Figure 2A). Transmission electron microscopy (TEM) revealed they have podovirus-like morphology, displaying polyhedral capsids with diameter of 94LJ±LJ3LJnm, tails with collar structures that were 34LJ± 3 nm with tail fibers with variable lengths (Figure 2B). From the calculated capsid size and genome length, we this phage packages its DNA at a density of 0.54 bp/nm^3^. This genome lacks direct terminal repeat sequences, stop-codon reassignment, and lysogeny-related genes (Table S1).

**Figure 2:**
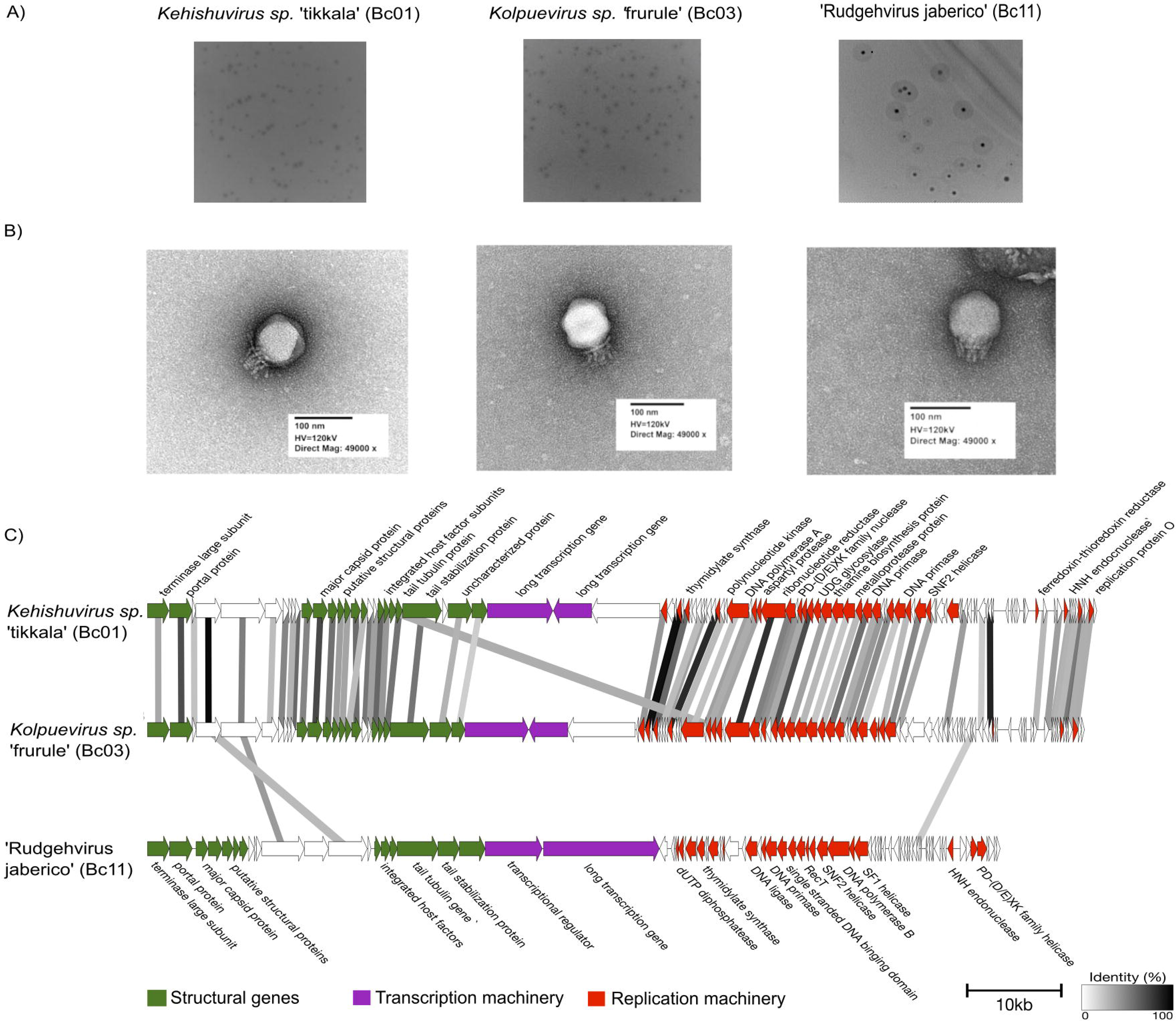
A) Plaque morphology of three species, ‘K. tikkala’ strain Bc01, ‘K. frurule’ strain Bc03, and ‘R. jaberico’ strain Bc11 B) Transmission electron microscopy images negatively stained with uranyl acetate of the three isolates C) Gene arrangement and functional annotation of the three genomes color-coded based on their functional modules and hypothetical genes represented in white. The direction of the arrows represents the direction of the gene read from the genome, and the arrows themselves represent individual genes. The links connecting the genes indicate sequence identity, ranging from 30% (grey) to 100% (black).

**Table 2:** Genome characteristics of the three novel *Crassvirales* species.

| Genome | length<br>(bp) | GC<br>% | Coding<br>density | Number of<br>CDS | Unknown<br>function | tRNA | DTR |
| --- | --- | --- | --- | --- | --- | --- | --- |
| *Kehishuvirus*<br>sp. 'tikkala'<br>strain Bc01 | 100,841 | 35.09 | 91.84 | 104 | 58 | 24 | False |
| *Kolpuevirus*<br>sp. 'frurule'<br>strain Bc03 | 99,523 | 33.00 | 92.06 | 108 | 63 | 5 | False |
| 'Rudgehvirus<br>jaberico' strain<br>Bc11 | 90,575 | 29.15 | 87.45 | 84 | 48 | 0 | False |

*Kolpuevirus sp.* ‘frurule’ strain Bc03 shares the *Steigviridae* family with ‘K. tikkala’ strain Bc01. This genome is 99,523 bp long with GC content of 33 %, 108 genes and four tRNA genes encoding arginine, asparagine, and tyrosine (Table 2). Similar to ‘K. tikkala’ strain Bc01, this phage also formed clear, uniform circular spot plaques but forming 2.3*10^9^ PFU/mL (Figure 2A). Displaying a podovirus morphology, the virion was slightly larger than K. tikkala’ strain Bc01 with capsids of diameter 97 ± 3 nm, a tail with collar structures of 33 ± 3 nm (Figure 2B), packaging its DNA at a lower density of 0.48 bp/nm^3^. Annotation of genes confirmed absence of direct terminal repeats, stop-codon reassignments, and lysogeny-related genes (Table S2). This is the first isolate within its genus.

‘Rudgehvirus jaberico’ strain Bc11 belongs to *Intestiviridae* family in a novel genus. This genome is 90,575 bp long with 29.15% GC content, encoding 84 genes and lacks tRNA genes (Table 2). Unlike the above two species, this isolate formed plaques with a circular halo, indicating depolymerase activity, forming 3.75 x10^3^ PFU/mL (Figure 2A). This isolate’s virion was relatively smaller in size compared to the other two isolates, with tails measuring 25 ± 4 nm (Figure 2B). Despite the smaller capsid size and genome, this phage packages its DNA at a density of 0.56 bp/nm^3^, comparable to ‘K. tikkala’ strain Bc01. Similar to the other two genomes, direct terminal repeats, stop-codon reassignments, and lysogeny-related genes (Table S3) were absent.

Comparative analysis across the three isolates shows that ‘K. tikkala’ strain Bc01 and ‘K. frurule’ strain Bc03, belonging to the same family (Steigviridae) exhibit higher gene similarity with each other. In contrast, ‘R. jaberico’ strain Bc11 from Intestiviridae family displays distinct gene arrangements (Figure 2C). Notably, all three genomes share two structural genes. encoding tail spike proteins with a domain encoding for polysaccharide degrading enzymes such as glycoside hydrolase. Structural protein 1 encompassing the ‘K. tikkala’ strain Bc01 protein (WEU69744.1) shared 97 % sequence similarity with the ‘K. frurule’ strain Bc03 protein (WEY17522.1), while collectively these sequences share greater than 39 % similarity with ‘R. jaberico’ strain Bc11 protein (WEU69859.1) (Figure S2). Similarly, structural protein 2 encompassing ‘K. tikkala’ strain Bc01 protein (WEU69745.1) shared 59 % sequence similarity with ‘K. frurule’ strain Bc03 protein (WEY17523.1), and together share more than 46% similarity with ‘R. jaberico’ strain Bc11 protein (WEU69857.1) (Figure S3).

### Synteny across all seven *Crassvirales* species successfully isolated

The comparison of the three novel species from this study that infect the same bacterial host with the four isolate *Crassvirales* genomes that infect other *Bacteroides* hosts, showed expected gene similarity based on their taxonomic assignment. Among the *Steigviridae* genomes, ‘K. tikkala’ strain Bc01 was most similar to *K. primarius* sharing 76 of 106 genes and the two genomes from *Wulfhauvirus* genus (strains DAC15 and DAC17) shared 115 of the 121 genes with greater than 30 % similarity. ‘K. frurule’ strain Bc03 belonging to a unique genus, *Kolpuevirus* exhibited intermediate similarity, sharing 68 genes with *Kehishuvirus* and 71 genes with *Wulfhauvirus* genus (Figure 3A). Within the *Intestiviridae* family, ‘R. jaberico’ strain Bc11 was compared to the *J. secundus,* and they shared 37 genes, including 11 structural genes, three transcription genes, and 23 replication-related genes (Figure 3B).

**Figure 3.**
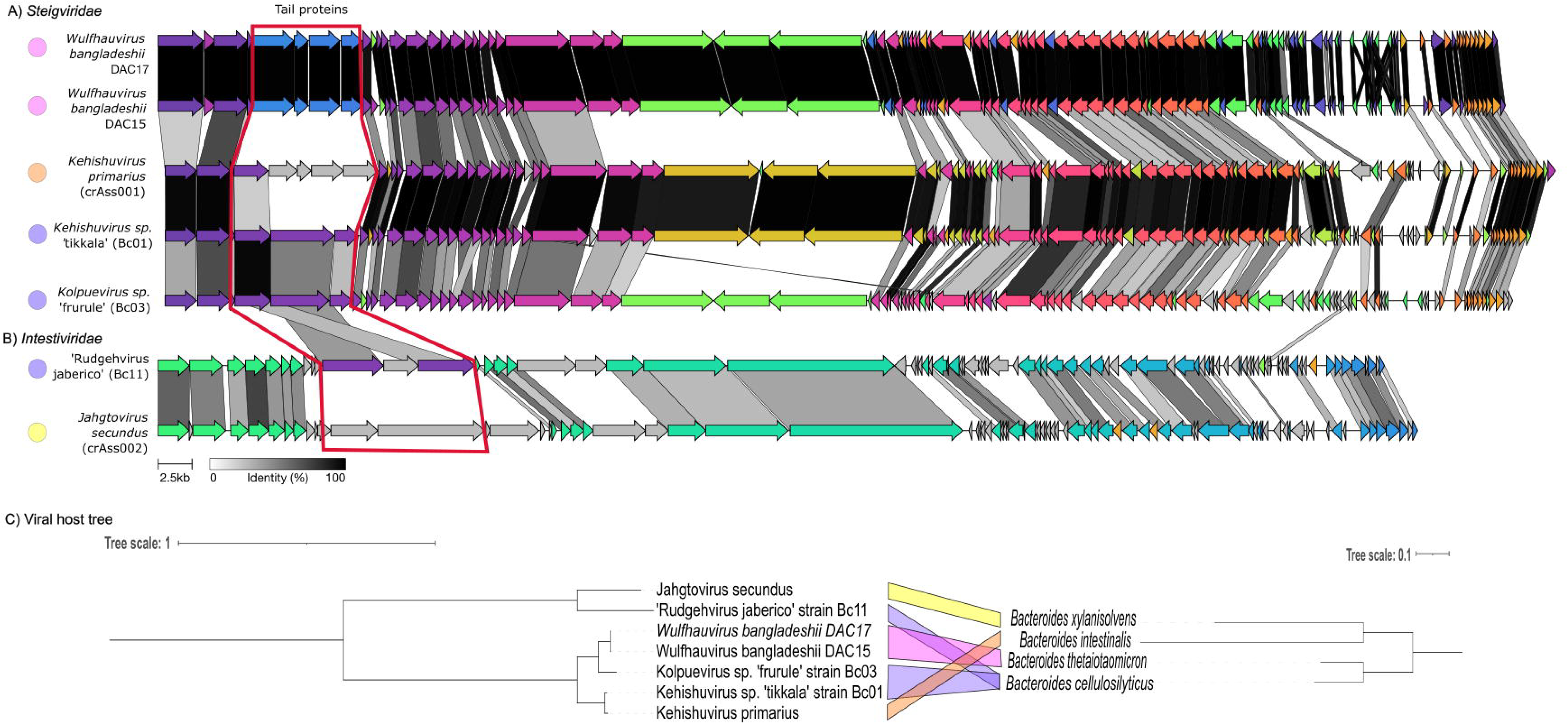
Gene synteny across seven pure culture isolates across two *Crassvirales* families A) *Steigviridae* family comprising five isolates spanning across three genera B) *Intestiviridae* family comprising two isolates from two genera. The arrows represent genes, with their direction indicating the gene direction, and their color indicating cluster group with the grey-colored arrows representing unique genes that didn’t form any clusters. Finally, the links connecting the genes are color-coded based on sequence similarity, ranging from grey (30%) to black (100%). The tail proteins that were shared between the three isolates from this study are highlighted with a red box. A dot is added next to each of the phage to represent the bacterial host, *B. thetaiotaomicron* in pink, *B. xylanisolvens* in orange, *B. cellulosilyticus* in purple, and *B. intestinalis* in pink C) Viral host-tree constructed using the portal gene for *Crassvirales* species and 16S gene for the bacterial hosts, with unique colors connecting the phage to its bacterial host.

The exception to the taxa-based similarity was the two structural genes, encoding tail spike proteins that were shared only among isolates infecting the same host, ‘K. tikkala’ strain Bc01, ‘K. frurule’ strain Bc03, ‘R. jaberico’ strain Bc11 despite belonging to different genera (Figure 2C).

As there were multiple *Crassvirales* species infecting multiple bacterial hosts (Figure 1A), we performed a coevolutionary test using Parafit (33) that supported random association between *Crassvirales* phages with their bacterial hosts (Parafit Global = 3.33, p values >0.05).

### Structural proteins playing a role in host interaction

To investigate the phage genes playing a role in host interaction, we compared all 1,887 genes across the 18 *Crassvirales* genomes, including 14 from this study that infect bacterial host, *B. cellulosilyticus* WH2 and the four *Crassvirales* isolates that infect four different bacterial hosts, *B. intestinalis, B. xylanisolvens,* and *B. thetaiotaomicron*. Together, from 18 *Crassvirales* isolates 1,766 genes were categorized into 383 orthologous groups (Table S4), while the remaining 121 genes remained singletons. To reinforce the validity of this analysis, we corroborated the species tree inferred from orthogroups (Figure S4) to have the same species level clustering as observed in phylogenetic tree (Figure 1A). There was one exception, *J. secundus* which belongs to *Intestiviridae* family was grouped with the *Steigviridae* isolates instead of its relative ‘R. jaberico’ strains, due to gene duplication or recombination events in this genomes.

Following the species level clustering, we identified 64 orthogroups (193 genes) that were specific to *Kehishuvirus,* 55 orthogroups (564 genes) specific to *Kolpuevirus,* 89 orthogroups (187 genes) specific to *Wulfhauvirus,* 73 orthogroups (148 genes) specific to ‘Rudgehvirus’, and 5 orthogroups (10 genes) specific to *Jahgtovirus* genera (Table S4). Within these groups, only two orthogroups–OG000000 (including Bc01: WEU69745.1, Bc03: WEY17523.1, Bc11: WEU69858.1) and OG000008 (including Bc01: WEU69744.1, Bc03: WEY17522.1, Bc11: WEU69859.1) included genes only from the 14 *Crassvirales* isolates that infect the same bacterial host, *B. cellulosilyticus* WH2. However, in orthogroup OG000000, four gene duplication events occurred with at least 50% of the descendant species having retained both the gene duplicates, therefore this orthogroup was not investigated further. Conversely, there were no gene duplication events within OG000008.

To determine if the genes in OG000008 is undergoing selection pressure, we calculated the number of synonymous (d_S_) and non-synonymous (d_N_) mutations occurring (d_N_/d_S_ <1). Averaging all the sequence pairs, we used the codon-based z-test to identify genes under selection and found that OG000008 rejected the null hypothesis (z-score=0.56, p-value<0.001), suggesting that these genes is under purifying selection. As recombination can impact this analysis, we ran Genetic Algorithm for Recombination Detection (GARD) to detect recombination, which identified five recombination breakpoints of which none were significant to be detected by the genetic algorithm. We therefore investigated the tail spike protein structure and role in host interaction.

### Tail spike protein interacts with TonB-dependent receptors

To identify the potential host interactions, we predicted structure of all 103 proteins from ‘K. tikkala’ strain Bc01, 109 proteins from ‘K. frurule’ strain Bc03, and 83 proteins ‘R. jaberico’ strain Bc11 were generated using Colabfold (34) (Protein structures available at doi.org/10.25451/flinders.21946034). Specifically, we compared the folded structures of tail spike proteins belonging to orthogroup OG000008 (Bc01: WEU69744.1, Bc03: WEY17522.1, Bc11: WEU69859.1) using Flexible structure AlignmenT by Chaining Aligned fragment pairs allowing Twists (FATCAT-rigid) method to show ‘K. tikkala’ strain Bc01 is similar to ‘K. frurule’ strain Bc03 with root-mean square deviation (RMSD) of 5.86 Å and only 74 % of paired residues in the structural alignment (Figure 4A). On the other hand, the tail spike protein of ‘K. tikkala’ strain Bc01 exhibited a RMSD of 6.61 Å and 60 % identity when compared to ‘R. jaberico’ strain Bc11 (Figure 4B).

**Figure 4:**
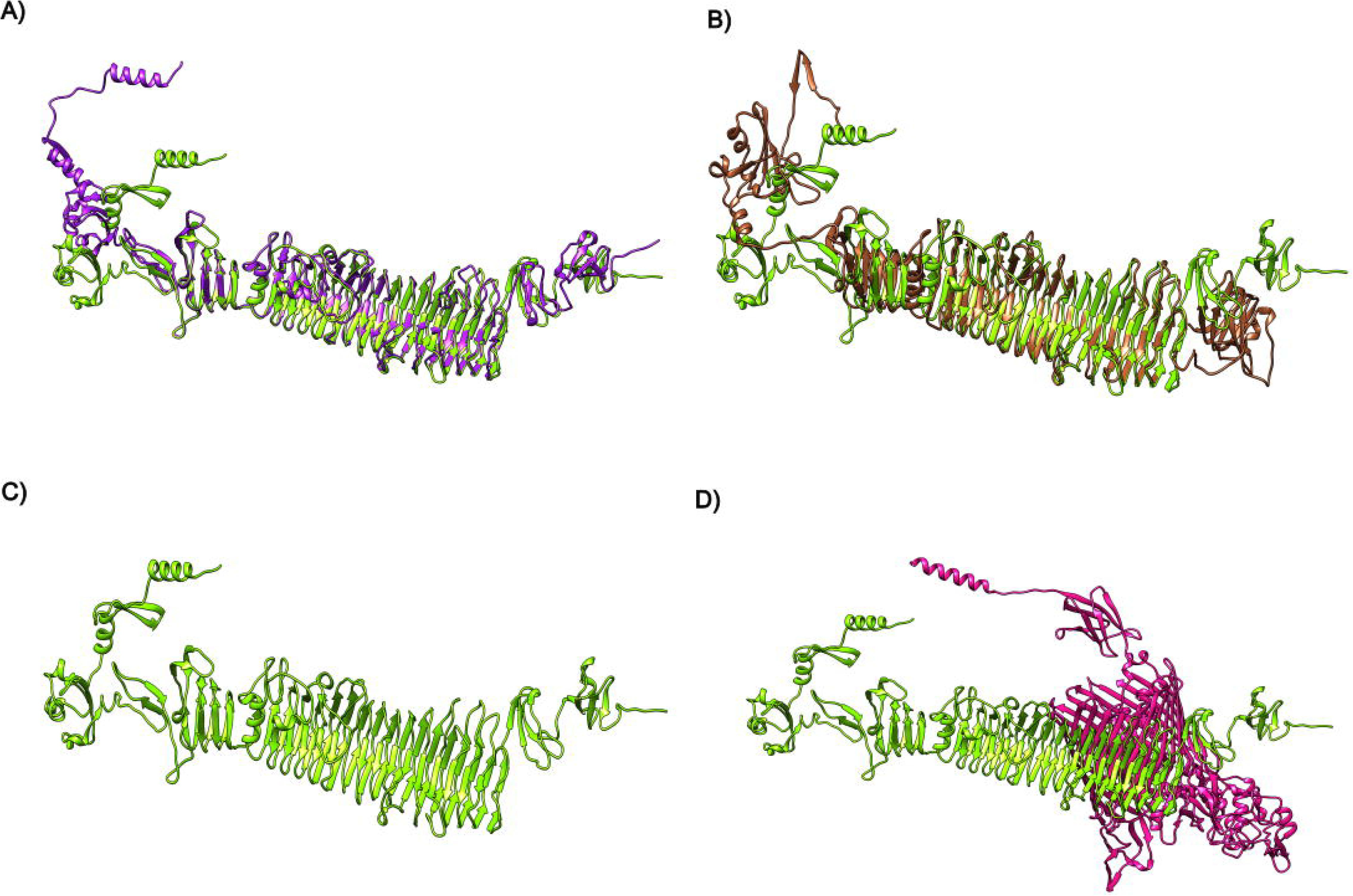
3D structure of tail spike proteins visualized using Chimera A) Structural alignment of tail spike protein *Kehishuvirus* sp. ‘tikkala’ strain Bc01 (WEU69744.1 in green) with *Kolpuevirus* sp, ‘frurule’ strain Bc03 (WEY17522.1 in purple). B) Structural alignment of tail spike protein *Kehishuvirus* sp. ‘tikkala’ strain Bc01 (WEU69744.1 in green) with ‘Rudgehvirus jaberico’ strain Bc11 (WEU69859.1 in brown). C) 3D Structure of *Kehishuvirus* sp. ‘tikkala’ strain Bc01 (WEU69744.1 in green) D) *Kehishuvirus* sp. ‘tikkala’ strain Bc01 (WEU69744.1 in green) docked with *Bacteroides cellulosilyticus* WH2 TonB-dependent receptor (A0A0P0GGA2 in pink).

Each of the tail spike protein structures was individually docked against all 3,223 predictions from the *B. cellulosilyticus* WH2 proteome available in the AlphaFold database using hdock-lite (Table S5). ‘K. tikkala’ strain Bc01 tail spike protein (WEU69744.1) (Figure 4C) interacted best with TonB-dependent receptors (UniProt ID: A0A0P0GGA2, hdock-score =-700) (Figure 4D), ‘K. frurule’ strain Bc03 protein (WEY17522.1) with another TonB-dependent receptor (UniProt ID: A0A0P0GR14, hdock-score= -694), and ‘R. jaberico’ strain Bc11 protein (WEU69859.1) with a different TonB-dependent receptor (UniProt ID: A0A0P0FZA4, hdock-score= -574).

## Discussion

The role of *Crassvirales* genomes in the human gut is enigmatic, which has been hindered by the limited number of cultured *Crassvirales* phages. Here, we address this gap by successfully isolating three novel *Crassvirales* species infecting *Bacteroides cellulosilyticus* WH2, belonging to different genera and families. This observation confirmed that the phages are not co-evolving with their bacterial hosts, rather have a shared ability to exploit similar features in their bacterial host. Notably, we identified a unique tail spike protein shared among isolates infecting the same bacterial host, undergoing purifying selection and interacting with the TonB-dependent receptors on the bacterial surface.

The *Crassvirales* order is currently comprised of vast and diverse collection of genomes. Despite this, the study of these phages has been limited due to the scarcity of pure isolates. The challenge associated with successful isolation underscores the difficulty in identifying and predicting their associated bacterial hosts. In our study, we addressed this challenge through focusing on wastewater samples, a source for phages infecting different *Bacteroides* hosts. Employing this approach, we successfully isolated 14 novel *Crassvirales* isolates specifically infecting *B. cellulosilyticus* WH2. These isolates were sequenced on different sequencing platforms, including Oxford Nanopore, Illumina Miseq or a combination of both. Nanopore assemblies provided high-quality and complete assemblies, however required polishing the assembly with Illumina reads to correct for frameshift errors that can fragment genes (35–37). As a result, the 14 complete genomes were classified at the family and genus levels, denoted as three novel species (Figure 1A), while the remaining isolates were grouped as strains of the same species (Table 1). The highest confidence isolate was selected for each species, *Kehishuvirus* sp. ‘tikkala’ strain Bc01, *Kolpuevirus* sp, ‘frurule’ strain Bc03 and ‘Rudgehvirus jaberico’ strain Bc11 and examined further.

Taxonomic assignment of the three novel species showed they belong to two families. *Kehishuvirus* sp. ‘tikkala’ strain Bc01 and *Kolpuevirus* sp, ‘frurule’ strain Bc03 were assigned to the *Steigviridae* family. This family also comprised of three other *Crassvirales* phages, *Kehishuvirus primarius, Wulfhauvirus bangladeshii* DAC15, and *Wulfhauvirus bangladeshii* DAC17 infecting other *Bacteroides* hosts. Notably, the two novel isolates exhibited clear, uniform circular spot morphology distinct from the turbid plaques observed in *K. primarius*, despite their close relationship within the same family. The third novel species, ‘Rudgehvirus jaberico’ strain Bc11 belonged to *Intestivirdae* family, along with other *Crassvirales* species, *Jahgtovirus secundus.* ‘R. jaberico’ strain Bc11 presented plaques with a circular halo surrounding the cleared spot, indicating depolymerase activity to break down the polysaccharides found on the bacterial cell wall.

Furthermore, comparing the three novel species, we found that their virion production, estimated from the number of plaques formed, was correlated to the number of tRNA genes within the genome (38). However, it is possible there are other factors such as gene regulation, and host immune responses that could also be influencing virion production. Additionally, we conducted genome density analysis in association with capsid sizes and genome lengths, revealing an inconsistency with prior isolated *K. primarius* and *J. secundus* species. The capsid diameters of the new three novel *Crassvirales* species of virions (90 to 97 nm) were apparently 20 % larger in size than those reported for *K. primarius* and *J. secundus* virions (77 nm) (27,29). However, considering that the reported values for *K. primarius* and *J. secundus* corresponded to the inscribed rather than the circumscribed diameters, a geometric correction of 22% that brought the genome density near 0.5 bp/nm^3^. This correction aligned with a larger diameter measured in the recently published cryo-EM reconstruction of *K. primaries* (32). The finding highlights the importance of accurately assessing virion dimensions and genome density to ensure consistency in the classification of *Crassvirales* phages.

The addition of the three novel *Crassvirales* species spanning multiple families infecting one bacterial host *B. cellulosilyticus* WH2 indicated these species may not be co-evolving with their bacterial hosts. We therefore tested all the successfully cultured *Crassvirales* species and their respective bacterial hosts to discover that they do not exhibit co-evolutionary patterns, rather support random association. These findings imply that the phage-host association within *Crassvirales* group are shaped by the environment and host interactions (33,39). Additionally, genome comparison of the known *Crassvirales* species showed greater shared similarity within genera. However, the three *Crassvirales* species despite belonging to three different genera shared two unique structural genes. Evolutionary analysis confirmed one of the two structural genes, encoding tail spike protein (comprising Bc01: WEU69744.1, Bc03: WEY17522.1, Bc11: WEU69859.1) formed an orthologous group, and is undergoing purifying selection pressure. Tail spike proteins have been shown to play crucial for binding to specific membrane receptors on the bacteria in tailed bacteriophages (40). Therefore, through preserving this gene function, the phage can successfully infect and replicate within the host.

We found the tail spike proteins of the three novel *Crassvirales* species to interact with different TonB-dependent receptors on the bacterial surface, provide significant insights into the mechanism of phage-host interactions. The bacterial host, *B. cellulosilyticus* WH2 possesses a substantial repertoire of up to 112 TonB-receptors on its surface. These receptors are typically used by *Bacteroides* to take up starches (41), and have been associated with phage sensitivity (30,31). The tail spike protein also encodes for polysaccharide degrading enzymes such as glycoside hydrolase domain that target the capsular polysaccharides on the bacterial surface, allowing for phage-host interaction and lead to infection. This interaction therefore ensures successful propagation, highlighting the evolutionary adaptation between the *Crassvirales* phage and their bacterial hosts.

Overall, our study on the three novel *Crassvirales* species infecting *Bacteroides cellulosilyticus* WH2 revealed critical insights into their evolutionary dynamics and interactions with the bacterial host. The novel phages belonging to different genera but infect the same host provide a valuable model system for studying the interactions that occur within one of the dominant members of the gut microbiome.

## Methods

### Phage sampling

Untreated sewage water (influent) was collected from a waste treatment plant in Cardiff, CA in 1L Nalgene bottles. An aliquot of 30 ml influent was centrifuged at 5,000 RCF for 5 min to pellet the debris. The supernatant was decanted and passed through a 0.22 μm pore size Sterivex filter. The filtrate was used as a phage source and stored between 2 to 8 L.

### Host bacteria cultivation

Bacterial species, *B. cellulosilyticus* WH2 (42) received as glycerol stocks from Washington University, St. Louis, *B. fragilis* NCTC 9343 (ATCC 25285), *B. stercoris* CC31F (ATCC 43183), and *B. uniformis* ATCC 8492 were received as glycerol stocks from BEI resources were used as bacterial hosts. All the bacteria were grown in brain-heart infusion media supplemented with 2 mM MgSO_4,_ and 10 mM MgCl_2_ we denote as BHISMg. Culture plates were supplemented with 1.5 % w/v agar and incubated at 37 L for 48 hrs under anaerobic conditions with 5 % H_2_, 5 % CO_2,_ and 90 % N_2_. Following incubation, an isolated colony was transferred into a 12 hrs deoxygenated BHISMg broth. Following anaerobic incubation at 37 L for 24 hrs the liquid cultures were further sub-cultured into another BHISMg broth and incubated overnight.

### Plaque assays

BHISMg plates were deoxygenated for 12 hrs in the anaerobic chamber and pre-warmed before use. For top agar plates were prepared by adding cooled molten BHISMg with 0.7 % w/v agar was inoculated with 500 µl of bacteria, and between 2 µl and 50 µl of processed phage influent. The plates were cooled for 15 min before incubating at 37 °C for up to five days. Plates were assessed daily for the development of plaques.

### Lysate preparation

Plaque from each plate was inoculated into 200 µl of SM buffer and homogenized to diffuse the phage from the agar to the buffer. A 200 µl aliquot of the phage was added to *B. cellulosilyticus* WH2 bacteria in the log-growth phase and grown at 37 °C anaerobically, overnight. The tubes containing the bacteria and phage were manually shaken every 30 min for the first three hours of incubation. Post incubation, tubes were centrifuged at 4500 *x* g for 5 min, and the supernatant was collected and concentrated using a 50,000 kDa MWCO Vivaspin ultrafiltration unit (Sartorius). Phage lysate was stored at 4 °C.

### Phage titering enumeration

Phage titer were enumerated using the molten agar overlay method described above. A 200 µl aliquot of the lysate was diluted 10-fold in sterile SM buffer, and 10 µl was spotted onto a BHISMg plate. The plates were incubated for 24-48 hrs at 37 °C. After incubation, the plates were analyzed by counting the plaques obtained to determining the titer.

### Viral DNA extraction and sequencing

Phage DNA was extracted using a Phage DNA isolation kit (Norgen) as per manufacturer instructions. In short, 1 ml of phage lysate was DNase I treated, lysed, and treated with Proteinase K. The sample was added to a spin column and washed three times. DNA was eluted in 75 µl of the elution buffer. The second elution recommended by the kit was not performed. The DNA obtained was quantified using a Qubit 1x dsDNA high-sensitivity assay kit (Invitrogen, Life Technologies) per manufacturer’s instructions. Oxford Nanopore MinION sequencing was undertaken the manufacturer’s instructions. In short, a maximum of 400 ng of sample DNA was used for the library preparation using Oxford Nanopore Rapid Barcoding Sequencing Kit (SQK-RBK0004); samples were barcoded, pooled and cleaned. The pooled samples was loaded and run on a Flowcell R9.4.1 (FLO-MIN106) following the manufactures instructions. The Illumina sequencing libraries were prepared by extracting the total nucleic acid (RNA and DNA) using the COBAS AmpliPrep instrument (Roche), with NEBNext library construction and sequenced on Illumina MiSeq using the paired-end 2x250 bp protocol as described in (43). The sequencing data were deposited to Sequence Read Archive in Bioproject, PRJNA737576.

For the Nanopore sequenced isolates, basecalling was performed with Guppy v6.0.1 with model dna_r9.4.1_450bps_hac. The reads were then processed with Filtlong v.0.2.20 (44) to remove reads less than 1,000 bp in length and exclude 5% of the lowest-quality reads. Similarly, Illumina sequences were processed with prinseq++ v.0.20.4 (45), filtering reads less than 60 bp in length, reads with quality scores less than 25, and exact duplicates.

### Genome assembly

To assemble the genomes, a pipeline based on Snakemake using Snaketool (46) was employed. Nanopore reads were assembled using Flye v2.9 (47), while Illumina reads were assembled using MEGAHIT v1.2.9 (48). These assemblers were selected as they provide assembly graphs, which are useful for completing fragmented genome assemblies, (49–53).

To evaluate assembly quality, the resulting contigs were processed with ViralVerify v1.1 to detect viral contigs (54), read coverage was calculated using CoverM v0.6.1(55), and the assembly graph was examined. The assembly graph provides information on connecting unitigs (high-quality contigs) representing the longest non-branching paths joined together to form contigs.

From each assembly, unitigs meeting specific criteria were selected as complete phage genomes. These unitigs had a length greater than 90 kb, were identified as viral, exhibited the highest read coverage and were classified as complete using CheckV v1.0.1. To ensure representation, one unitg per sample was selected as the complete phage assembly. In the end, the assemblies were polished with high-coverage Illumina reads using Polca to reduce sequencing-related errors (56).

Among the 41 phage genomes, 14 phages infecting *B. cellulosilyticus* were identified as belonging to *Crassvirales* order. These genomes were approximately 90 to 120 kbp in length and aligned against known *Crassvirales* genomes. Among these phages, eight samples were sequenced on both Nanopore and Illumina sequencing platforms (Bc01 to Bc03, Bc05 to Bc08), while four were sequenced only on Nanopore platforms (Bc09 to Bc11), and the remaining four were sequenced only on Illumina platforms (Bc04, Bc12 to Bc14).

### Taxonomic and functional annotation

The isolates in this study were processed with CrassUS (57), a specialized tool for annotating *Crassvirales* genomes, providing taxonomic, functional annotation along with direct terminal repeats (DTR), and average nucleotide identities (ANI) of similar reference genomes. Taxonomic annotations from CrassUS followed ICTV (20,22) criteria for *Crassvirales* order demarcation. Phylogenetic trees were constructed using conserved genes (MCP, portal, *terL*) with MAFFT v7.49 (58) for alignment, followed by trimal v1.4.1 for trimming, and FastTree v.2.1.10 (59) for inference. Trees were built using JTT model, CAT approximation with 20 rate categories, and visualized using iTol (60).

To compare the predicted genes and their arrangement across species, clinker plots (61) were used after re-circularizing the genes to start at the *terL.* This allowed for the examination of synteny across genomes. Additionally, tRNA genes encoded by the phages, which evade host translation machinery, were predicted with tRNA-scanSE (62).

### Phage-host co-phylogenetic analysis

To determine if the phage co-evolve with bacterial hosts, we performed a cophylogenetic analysis using Parafit (33) via the ape R package. The distance matrix of the trimmed multiple sequence alignment (MSA) using MAFFT v.7.520 (58) and trimal v1.4.1 of the seven *Crassvirales* species (’K. tikkala’ Bc01, ‘K.frurule’ Bc03, ‘R.jaberico’ Bc11, *K. primarius, J. secundus, W. bangladeshii* DAC15, *W. bangladeshii* DAC17) portal gene, was generated using EMBOSS distmat v6.6.0. These steps were repeated for the associated bacterial hosts, *B. cellulosilyticus, B. intestinalis, B. thetaiotomicron* and *B. xylanisolvens.* The two distance matrices were compared using Parafit (33) with 1000 permutation, and Cailliez eigen value correction. The trimmed MSA was used to generate the two phylogenetic trees using FastTree v.2.1.10 using JTT model, CAT approximation with 20 rate categories, and visualized using iTol (60).

### Transmission electron microscopy imaging

*Crassvirales* phages were grown using the phage overlay method described above. To prepare the phage lysates, they were diluted 1:10, and 5 μL of the diluted phage lysate was applied to a plasma-cleaned grid for two minutes at room temperature. The grids used were formvar and carbon coated 200 mesh grids and they were plasma cleaned using the Gatan (Solarus) Advanced plasma system for 30 sec prior to use. The excess phage lysate sample was wicked off with Whatman filter paper and the grid was washed with 5 μL of water. The sample was negatively stained with 5 μL of the 2 % w/v uranyl acetate for 1 minute. The excess stain was wicked off with filter paper to dry the sample on the grid. The grid was then imaged using a Tecnai G2 Spirit TEM operated at 120kV at a magnification of 49,000x and the images were recorded on an AMT Nanosprint 15 digital camera using software v7.0.1.

Phage measurements were conducted using the ImageJ software (63). The capsid diameter was calculated by measuring the diameter of the circle circumscribing the capsid, such that the more distant vertices of the projected capsid contacted the circle (Figure S5). The length of the tail was calculated from the base of the capsid to the end of the visible tail, including the collar section of the phage structure. Tail fibres or appendages was calculated (Figure S5). Average measurements from 5 phages were calculated and reported. The TEM image was further edited for publication using the GNU Image Manipulation Program (GIMP) (64).

The packing genome density was predicted by correcting the measured radius from the expected capsid thickness and calculating the internal volume assuming an icosahedral model inferred from prior tailed phage capsid studies (65,66). The results can be reproduced using the Colab notebook available at link, https://shorturl.at/enAKS.

### Evolutionary analyses

The 14 *Crassvirales* isolated and assembled genomes in this study and the four reference pure culture isolates, *K. primarius*, *J. secundus, W. bangladeshii* DAC15 and DAC17 were assessed together for this analysis. Orthologous genes were identified from genes predicted from the above 18 genomes, using Orthofinder v2.5.4 default settings to determine signatures for host interactions. The default settings use diamond for sequence search, MAFFT for alignment, FastTree for tree inference and STAG species tree method (67). Orthogroups that included genes present only in phages from the host *B. cellulosilyticus* WH2 were examined further.

These orthogroups were aligned using Muscle (68) codon-based multiple sequence alignment in MEGA11 (69). To test for codon-based positive selection, we calculated the probability of rejecting the null hypothesis of strict neutrality (d_N_ = d_S_), and in favour of the alternate hypothesis (d_N_ > d_S_). The d_N_/d_S_ values were calculated from the MSA using MEGA v.11.0 (70), with the Li-Wu-Luo method (71). The variance of the difference was computed using bootstraps, set to 100 replicates. As this analysis can be misleading in the presence of recombination breakpoints, orthogroups were run through Genetic Algorithm for Recombination Detection (GARD) analysis (72), with default settings. This method utilizes a combination of phylogenetic and statistical approaches to detect recombination signals.

### Predicting proteins 3D structure and docking

The 3D structures of the proteins from ‘K. tikkala’ strain Bc01, ‘K. frurule’ strain Bc03, and ‘R. jaberico’ strain Bc11 were predicted using Colabfold version 1.4.0 (34) on the Gadi server at the National Computational Infrastructure (NCI). To determine structural similarity, the protein structures were run through pairwise structure alignment using the Flexible structure AlignmenT by Chaining Aligned fragment pairs allowing Twists (FATCAT) that allows for flexible protein structure comparison (73,74).

To predict the phage protein interaction with the bacterial host, the previously predicted 3D protein structures of all the proteins for *B. cellulosilyticus* WH2 were downloaded from the AlphaFold Protein Structure Database via the Google Cloud Platform (75). All protein pairs were docked using hdock-lite version 1.1 (76) on the Gadi server. The results from hdock were sorted based on the binding score (hdock-scores) in the output file to identify the highest-quality binding predictions for each phage protein. In general, lower HDock-scores are indicative of more favorable or stronger protein-protein interactions, suggesting a higher likelihood of a stable complex formation. Higher HDock-scores, on the other hand, may suggest weaker or less favorable interactions.

The 3D structure of the proteins was visualized using Chimera.

## Author Contributions

BP performed bioinformatics analysis and wrote the paper. AAV, CS, SKG, MA, NJ, LB, CH and WSP collected samples, isolated and cultured phages. MFM, MA, KP and LD sequenced phages. CL and SKG took TEM images. KM, MJR, PD, SRG, VM, GB and AL performed bioinformatics analysis. SH, DW, AMS, EAD and RAE conceived the project, performed the bioinformatics, and wrote the paper with input from all authors.

## Supporting information

Bioinformatics workflow supplement

## Acknowledgments

This research/project was undertaken with the assistance of resources and services from Flinders University and the National Computational Infrastructure (NCI), which is supported by the Australian Government.

## Supplementary Figures

Figure S1: Showing the taxa classification of the three novel species remains consistent across the three conserved proteins A) portal gene, B) Major capsid protein (MCP), and C) terminase large subunit (terL). The outgroup across all three trees set to *Cellulophaga* phage phi13:2. The placement of the three novel species are highlighted on the tree, Bc01 belonging to *Kehishuvirus* genera (light green), Bc03 belonging to *Kolpuevirus* genera (purple), and Bc11 belonging to a novel genus named ‘Rudgehvirus’ (brown).

Figure S2: Multiple sequence alignment of shared structural protein 1 from Fig 2C including K. tikkala’ strain Bc01 protein, WEU69744.1, ‘K. frurule’ strain Bc03 protein, WEY17522.1, and ‘R. jaberico’ strain Bc11 WEU69859.1 (reference sequence), showcasing the sequence identity with amino acids that were not shared in grey, and the rest in different color, based on the amino acid group.

Figure S3: Multiple sequence alignment of shared structural protein 2 from Fig 2C including ‘K. tikkala’ strain Bc01 protein, WEU69745.1, ‘K. frurule’ strain Bc03 WEY17523.1, and ‘R. jaberico’ strain Bc11 WEU69857.1 (reference sequence), showcasing the sequence identity with amino acids that were not shared in grey, and the rest in different color, based on the amino acid group.

Figure S4: Species tree inferred from orthogroups using OrthoFinder and rooted to ‘Rudgehvirus jaberico’ strain Bc13 using the STRIDE algorithm. The 14 *Crassvirales* isolates are color coded based on their species classification, ‘Rudgehvirus jaberico’ strains in brown, *Kehishuvirus* sp. ‘tikkala’ strains in dark blue, and *Kolpuevirus* sp. ‘frurule’ strains in light blue. The four *Crassvirales* genomes from other studies are color coded in black. Further, to denote the family level classification, we added red dots next to *Intestiviridae* family members, and cyan dots next to *Steigviridae* family members.

Figure S5: TEM phage measurements were taken for A) Capsid diameter, by drawing a circle around the polygon with the edges within the circle. The diameter of this circle was measured and represented as the capsid diameter. B) For tail length, a line was drawn from the base of the capsid to the visible edge of the tail fibers. This was repeated over five phages of the same sample and an average with standard deviation was calculated across all of them.

## Supplementary Tables

Table S1: *Kehishuvirus sp.* ‘tikkala’ strain Bc01 functional annotation from crassUS

Table S2: *Kolpuevirus ‘*frurule’ strain Bc03 functional annotation from crassUS

Table S3: ‘Rudgehvirus jaberico’ strain Bc11 functional annotation from crassUS

Table S4: Orthologous groups identified across the 18 *Crassvirales* isolates, highlighted the two orthogroups that are present within the 14 *Crassvirales* isolates from this study, infecting the same bacterial host.

Table S5: Top 10 *Bacteroides cellulosilyticus WH2* proteins docked to *Kehishuvirus* sp ’tikkala’ strain Bc01 WEU69744.1, *Kolpuevirus* sp ’frurule’ strain Bc03 WEY17522.1, and ’Rudgehvirus jaberico’ strain Bc11 WEU69859.1 tail spike protein using hdock-lite.

