## Supplementary figures and images for "Host interactions of novel *Crassvirales* species belonging to multiple families infecting bacterial host, *Bacteroides cellulosilyticus* WH2"

### Bioinformatics workflow supplement

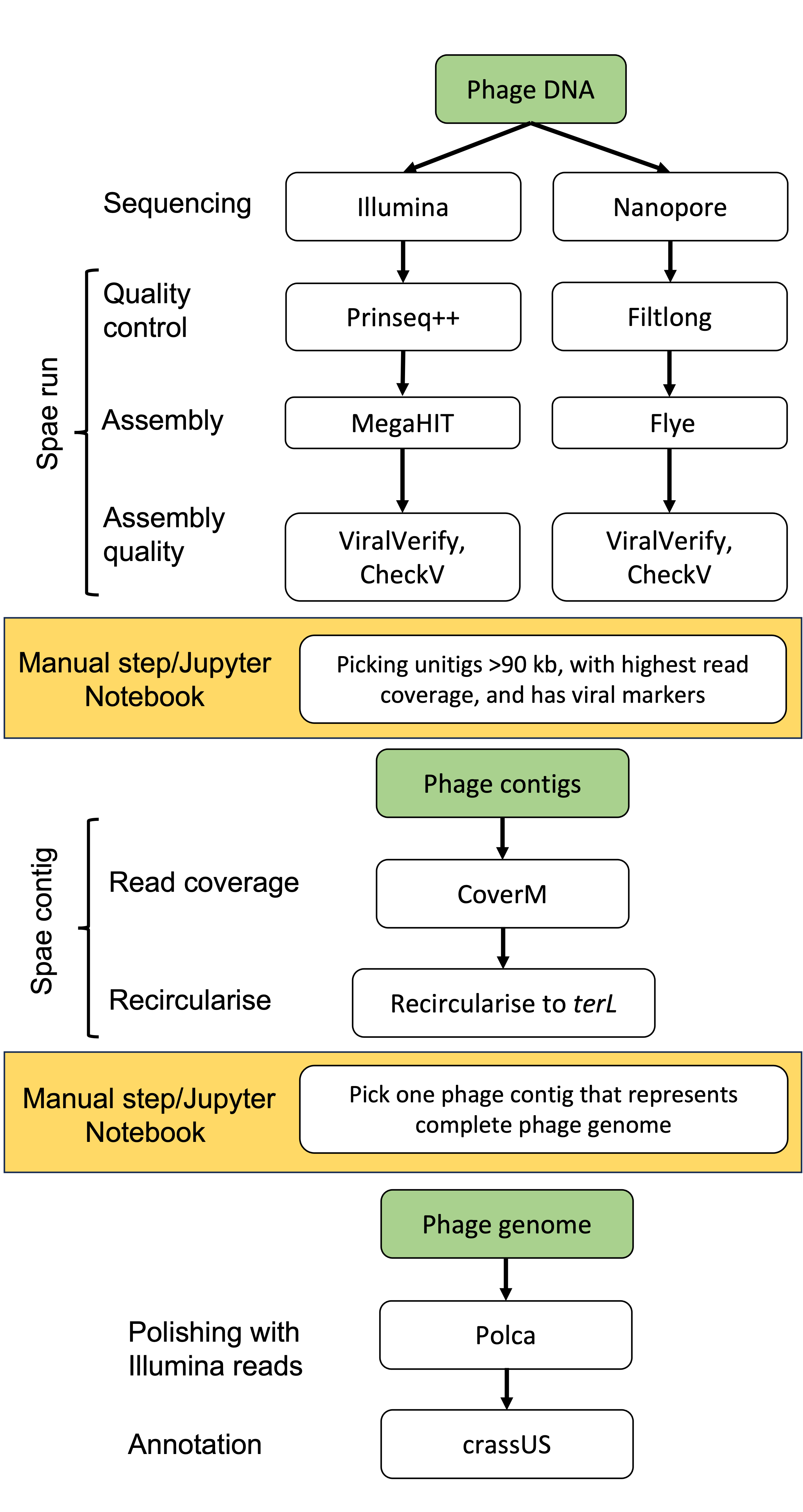

### Figure S1

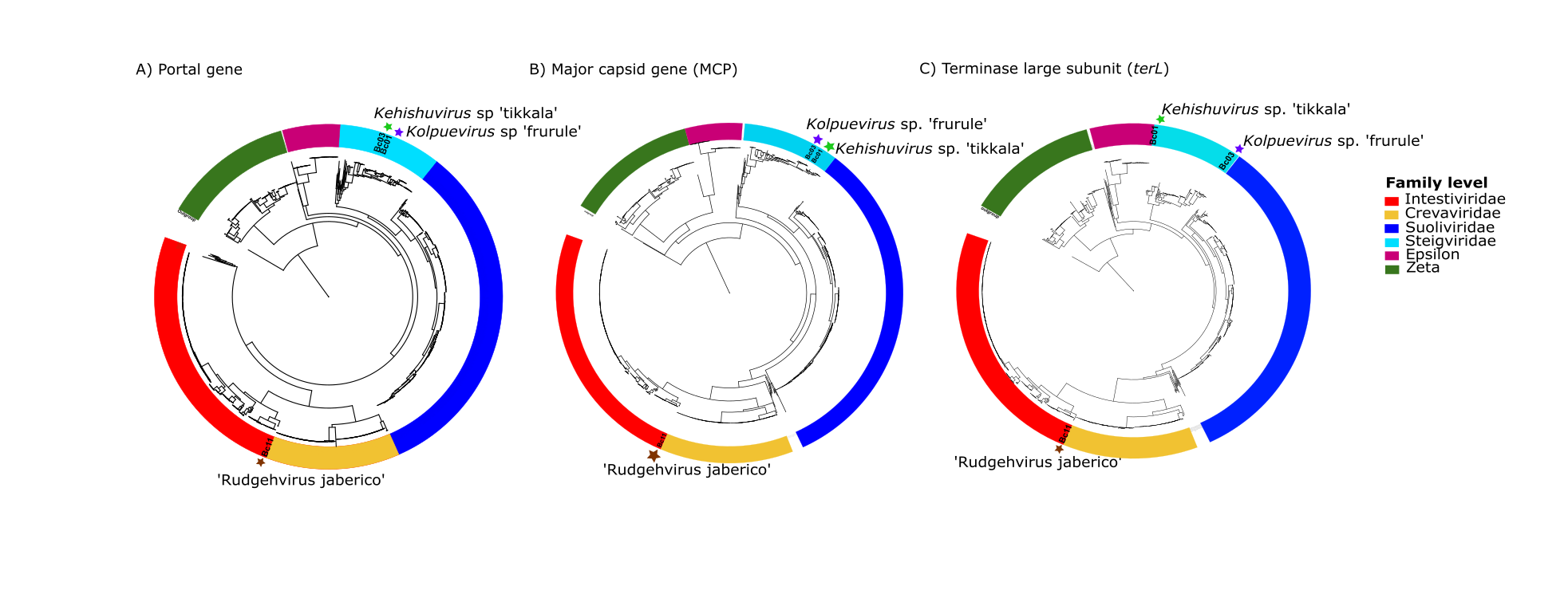

### Figure S2

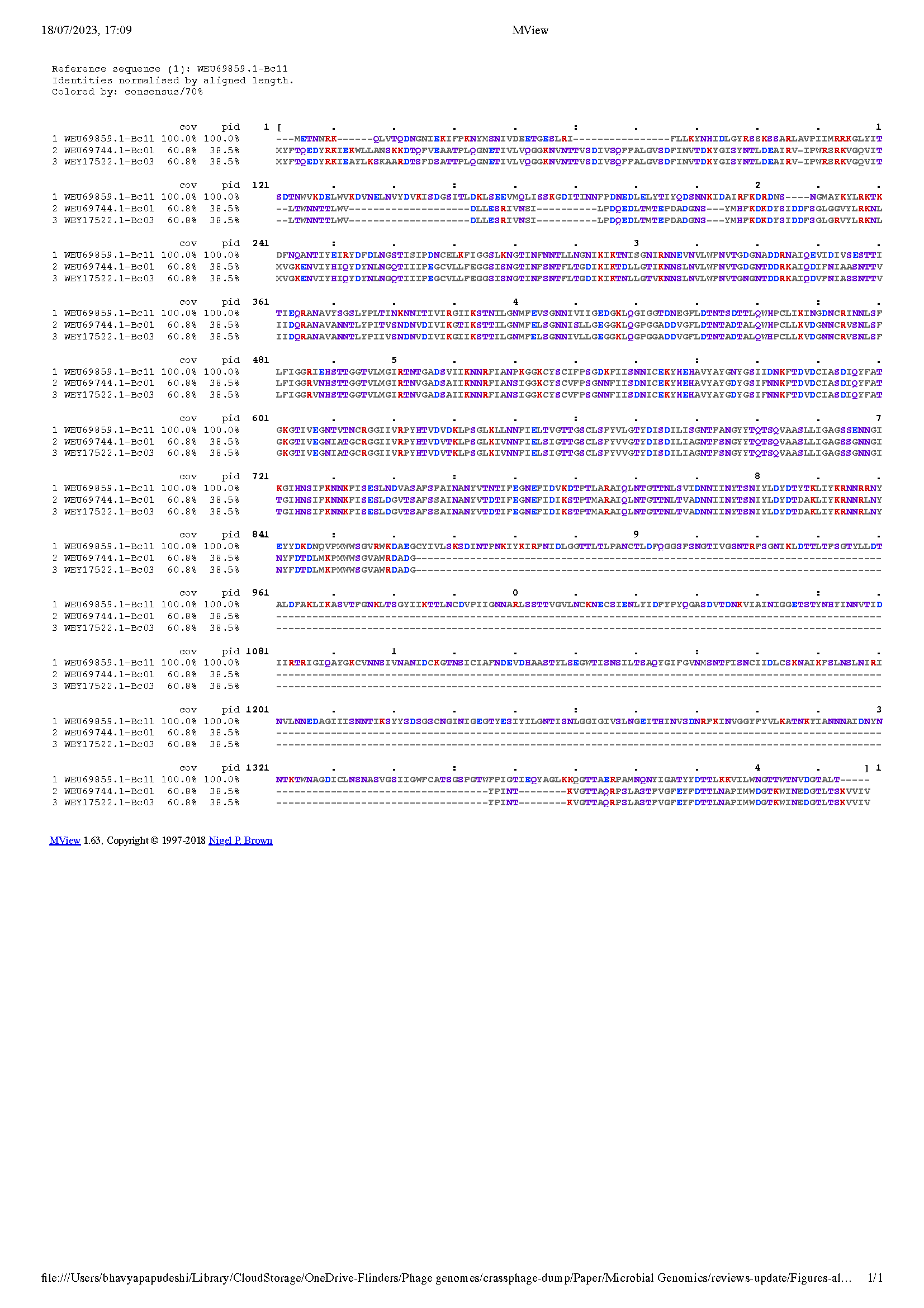

### Figure S3

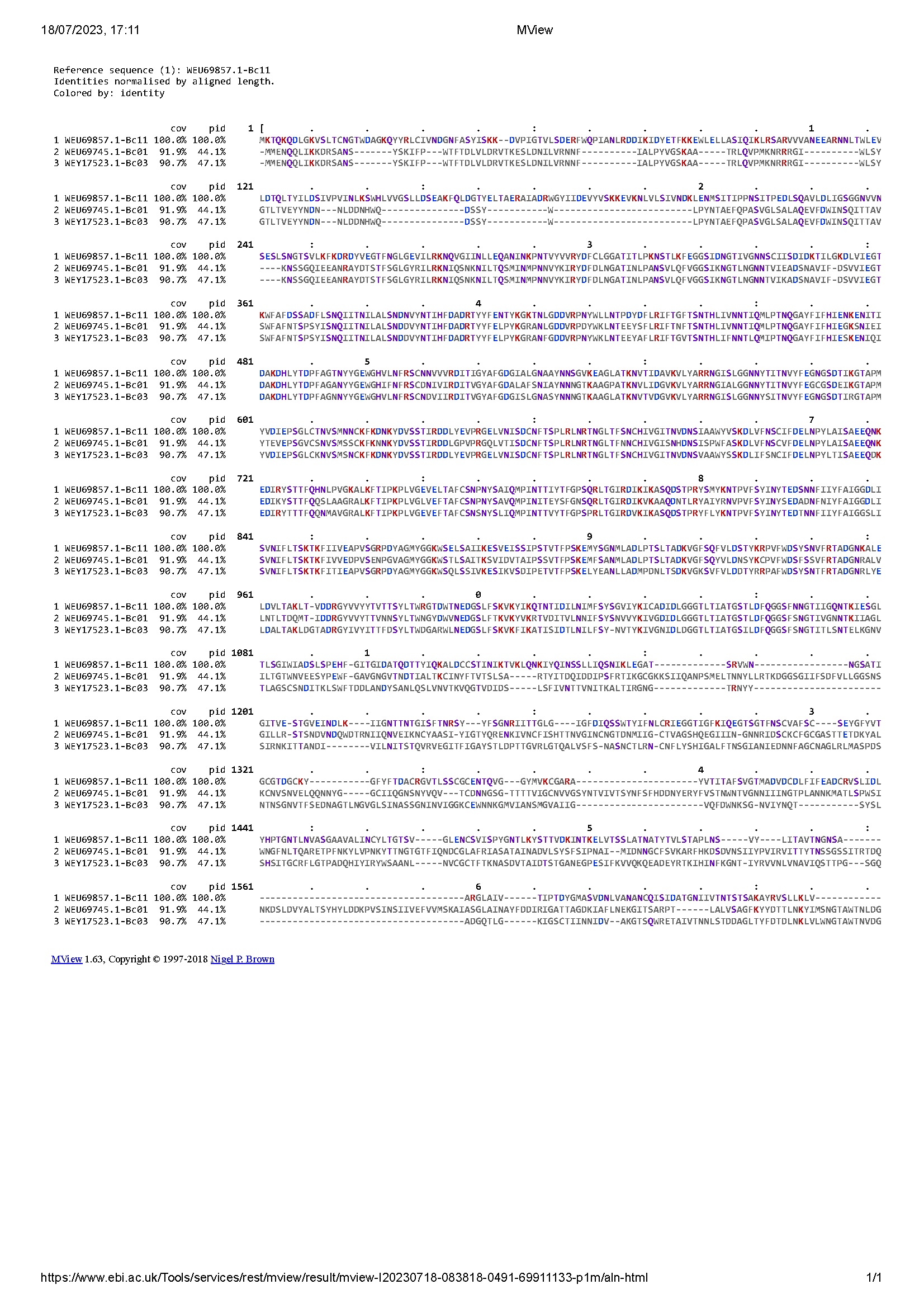

### Figure S4

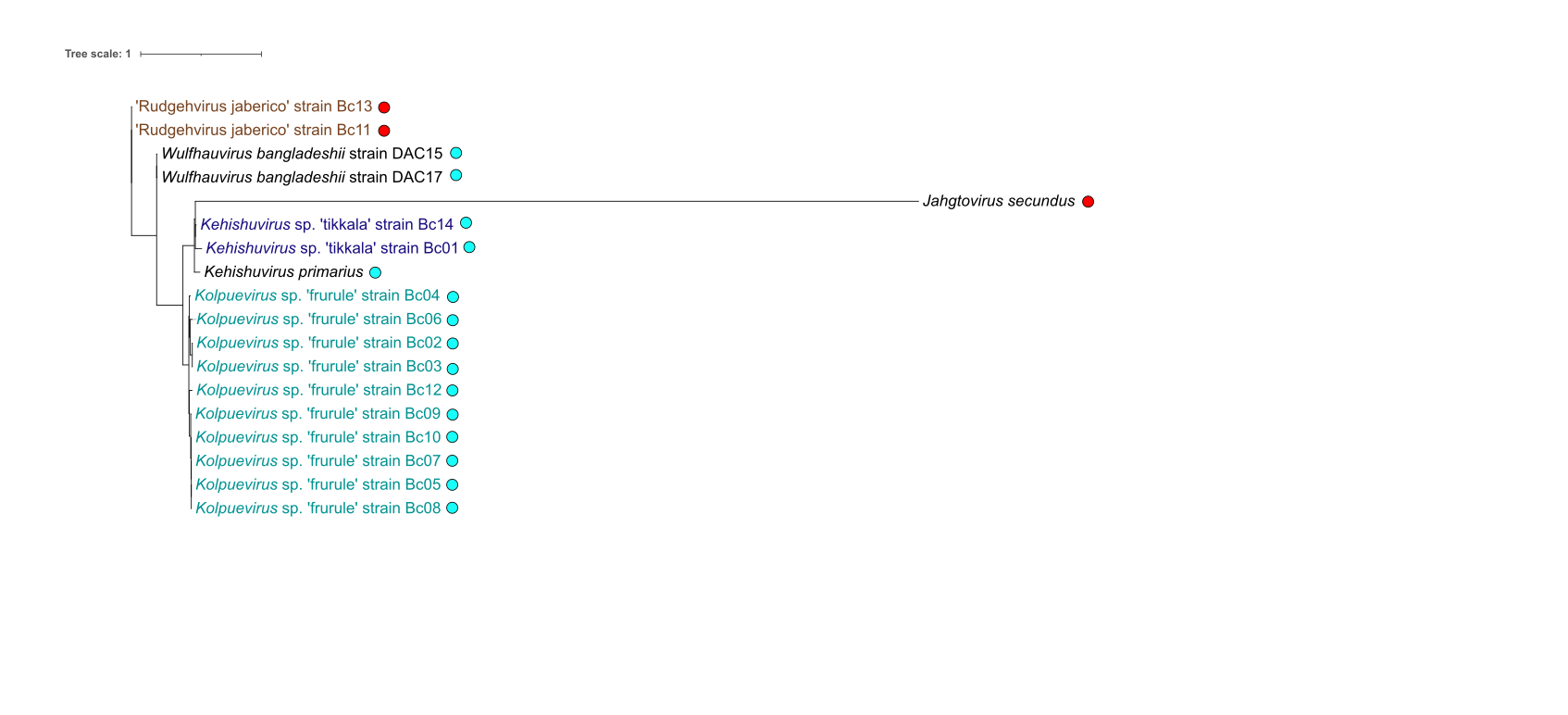

### Figure S5

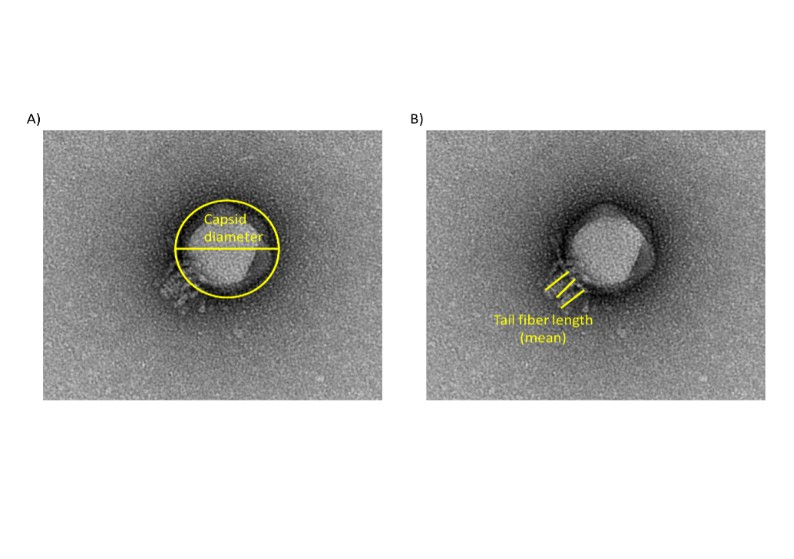
